# *In vivo* visualization of pig vagus nerve ‘vagotopy’ using ultrasound

**DOI:** 10.1101/2020.12.24.424256

**Authors:** Megan L. Settell, Maïsha Kasole, Aaron C. Skubal, Bruce E. Knudsen, Evan N. Nicolai, Chengwu Huang, Chenyun Zhou, James K. Trevathan, Aniruddha Upadhye, Chaitanya Kolluru, Andrew J. Shoffstall, Justin C. Williams, Aaron J. Suminski, Warren M. Grill, Nicole A. Pelot, Shigao Chen, Kip A. Ludwig

## Abstract

**Background:** Placement of the clinical vagus nerve stimulating cuff is a standard surgical procedure based on anatomical landmarks, with limited patient specificity in terms of fascicular organization or vagal anatomy. As such, the therapeutic effects are generally limited by unwanted side effects of neck muscle contractions, demonstrated by previous studies to result from stimulation of 1) motor fibers near the cuff in the superior laryngeal and 2) motor fibers within the cuff projecting to the recurrent laryngeal.

**Objective:** The use of patient-specific visualization of vagus nerve fascicular organization could better inform clinical cuff placement and improve clinical outcomes.

**Methods:** The viability of ultrasound, with the transducer in the surgical pocket, to visualize vagus nerve fascicular organization (i.e. vagotopy) was characterized in a pig model. Ultrasound images were matched to post-mortem histology to confirm the utility of ultrasound in identifying fascicular organization.

**Results:** High-resolution ultrasound accurately depicted the vagotopy of the pig vagus nerve intra-operatively, as confirmed via histology. The stereotypical pseudo-unipolar cell body aggregation at the nodose ganglion was identifiable, and these sensory afferent fascicular bundles were traced down the length of the vagus nerve. Additionally, the superior and recurrent laryngeal nerves were identified via ultrasound.

**Conclusions:** Intraoperative visualization of vagotopy and surrounding nerves using ultrasound is a novel approach to optimize stimulating cuff placement, avoid unwanted activation of motor nerve fibers implicated in off-target effects, and seed patient-specific models of vagal fiber activation to improve patient outcomes.

## Introduction

The therapeutic effects of vagus nerve stimulation (VNS) for epilepsy and heart failure, while significant in some patients, are often limited by intolerable side effects including throat tightening or pain, voice changes, hoarseness, cough, and dyspnea (Howland, 2014; Morris & Mueller, 1999). The inadvertent stimulation of somatic nerve branches extending from the vagus, such as the superior and recurrent laryngeal nerve (SLN and RLN, respectively), has been implicated as the cause of these side effects (Nicolai et al., 2020; Tosato et al., 2007; Yoo et al., 2013). These nerve branches are either activated through stimulation of fascicles within the stimulating cuff (RLN), or by current escaping the cuff (SLN) (Boon et al., 2009; Castoro et al., 2011; Nicolai et al., 2020). The SLN and RLN innervate neck muscles involved in many of the therapy-limiting side effects and therefore avoiding stimulation of these nerve fibers is paramount.

The vagus nerve (VN) contains a topographical organization (Settell et al., 2020), or vagotopy, that has the potential to be visualized using ultrasound. Previous work in a pig model of VNS demonstrated a bimodal organization in the VN. In the nodose ganglion pseudo-unipolar cell bodies (predominately sensory afferents) are grouped into a large fascicle, and distinct from a separate, smaller grouping of nerve fibers. This secondary grouping of nerve fibers give rise to the superior and recurrent laryngeal nerve branches (Settell et al., 2020). This bimodal arrangement of fascicles could be used to strategically place VNS cuffs to avoid the neuronal projections that innervate muscles implicated in side effects. Current clinical VNS stimulating cuffs wrap approximately 270° around the vagus nerve, and thus stimulates the circumference of the trunk mostly indiscriminately. Strategic placement of small electrodes and utilization of a current steering stimulation protocol, to target sensory over motor regions, could minimize therapy limiting activation of the neck muscles and optimize clinical efficacy.

Visualization of peripheral nerves using ultrasound could be an effective intraoperative method to identify fascicular organization and pertinent anatomical information *in vivo*. Ultrasound offers higher resolution, and is more cost-effective than other imaging modalities such as magnetic resonance imaging (MRI) (Zaidman et al., 2013). The use of ultrasound for neuropathology was first reported in the 1980s, with improvements in capabilities over the last thirty years (Cartwright et al., 2017). Non-invasive ultrasound has been completed in patients on a variety of superficial nerves demonstrating fascicular resolution. The sciatic nerve has been visualized in patients using ultrasound during popliteal sciatic nerve block for hallux valgus surgery (bunionectomy), with clear visualization of the epineurium through the skin (Karmakar et al., 2013). The median nerve (4 cm skin to nerve depth, 10 MHz transducer) (Marciniak et al., 2013), radial and ulnar nerves, are more superficial than the sciatic nerve and can be visualized through the skin during carpal tunnel evaluation with slightly better resolution of fascicles (Marciniak et al., 2013; Taylor et al., 2016).

Despite the ability to visualize these superficial nerves, visualizing fascicular organization of the VN with ultrasound poses a unique problem, as it is below skin, fat, and muscle. Current capabilities of the clinical transducers do not allow for high-resolution, non-invasive visualization of the fascicular organization of deep nerves such as the VN (Brown et al., 2016; Inamura et al., 2017). Though non-invasive ultrasound of the VN has been established in the clinical setting for diagnosis of masses of the neck (Giovagnorio & Martinoli, 2001), the depth of penetration is not sufficient to observe fascicular organization, and resolution tends to be poor (Inamura et al., 2017). In humans, the VN is 36.2□±□9.4□mm (mean ± SD) from the surface of the skin, with no differences between sides or sexes (Hammer et al., 2018). Given the depth of the VN, we propose a novel approach for visualizing vagotopy by placing the ultrasound transducer within the surgical pocket.

We demonstrate a novel intraoperative methodology for visualization of the vagotopy of the pig VN using a high frequency (50 MHz) ultrasound transducer within the surgical pocket. We aimed to use ultrasound to 1) identify the bimodal organization between the pseudo-unipolar cell bodies (sensory afferents) and the secondary fascicle grouping giving rise to the SLN and RLN at the level of the nodose ganglion, 2) resolve the bimodal organization of fascicles each of these groupings become within the surgical window, and 3) obtain additional information about the fasicular organization of the SLN and RLN themselves that may be useful in seeding computational models to inform off-target activation. Here, ultrasound images were matched to histological cross sections to confirm our real-time ultrasound identification of fascicular organization. In the future, real-time ultrasound can be collected, analyzed, and used to inform electrode cuff placement. This approach could lead to patientspecific, optimized placement of the stimulating cuff, resulting in reduced effects on off-target fibers and potentially more efficacious stimulation.

## Materials and Methods

### Subjects

All study procedures were approved by the Mayo Clinic Institutional Animal Care and Use Committee, and procedures were conducted under the guidelines of the American Association for Laboratory Animal Science in accordance with the National Institutes of Health Guidelines for Animal Research (Guide for the Care and Use of Laboratory Animals). Subjects included 3 healthy domestic (Yorkshire/Landrace crossbreed) swine (1F/2M; mean ± SD = 37.1 ± 5.11 kg). All subjects were housed individually (21°C and 45% humidity) with ad libitum access to water and were fed twice a day. Each subject was given an intramuscular injectable induction anesthesia: telazol (6 mg/kg), xylazine (2 mg/kg), and glycopyrrolate (0.006 mg/kg). An intramuscular injection of buprenorphine was given as an analgesic (0.03 mg/kg). Following induction, subjects were endotracheally intubated and maintained with a mechanical ventilator using 1.5-3% isofluorane. A blood pressure catheter was placed in the femoral artery (Millar, Inc., Houston, TX, Model # SPR-350S), and an intravenous catheter placed in the peripheral ear vein for drug and fluid administration. Subjects were endotracheally intubated and maintained with a mechanical ventilator using 1.5-3% isoflurane. All vital signs including temperature, heart rate, CO_2_, and respiration were continuously collected and recorded every 15 minutes and used to monitor depth of anesthesia.

### Surgical Methods

The surgical approach for exposing the VN and microdissection procedures have been described previously (Settell et al., 2020). Briefly, in a dorsal recumbence position, a ventral incision was made on the subject’s right side, just lateral and parallel to midline starting at the level of the mandible. Tissue was divided to locate the carotid sheath which was incised to expose the carotid artery, internal jugular vein, and VN. The VN was bluntly dissected from the nodose ganglion to approximately 10 cm caudal; careful measures were taken to avoid disturbing any of the surrounding branches, such as the SL or sympathetic trunk (ST). This exposed region spans the equivalent location for cervical VNS implantation in a patient, as identified by a practicing neurosurgeon (Nicolai et al., 2020; Settell et al., 2020). The incision site was kept moist with 0.9% sterile saline until the completion of experiment.

### Ultrasound

The ultrasound approach for this study was described previously (Huang et al., 2019). Briefly, after the surgical procedure, all ultrasound images were collected using a Vevo^®^ 3100 high frequency imaging system (FUJIFILM VisualSonics Inc., Toronto, Canada). The high frequency 50 MHz linear array transducer (MX700, 35 μm nominal axial resolution, 70 μm nominal lateral resolution) was placed within the surgical pocket (Figure 1A), 1-2 mm above the VN (Figure 1B). The surgical pocket was filled with mineral oil to increase coupling between the transducer and nerve, and the vagus nerve suspended from surrounding tissue using vessel loops to limit movement artifact and improve image quality. The transducer was attached to a linear stepper motor (P/N 11484, VisualSonics Inc.) connected to the Vevo^®^ integrated rail system to allow for smooth acquisition of images along the length of the nerve, without the need for manual manipulation. The transducer was directed to move along the length of the VN in the cranial to caudal direction, starting at the nodose ganglion and extending the length of the surgical window as 3D plane-by-plane volumetric B-mode images were collected (Figure 1C and 1D).

**Figure 1:**
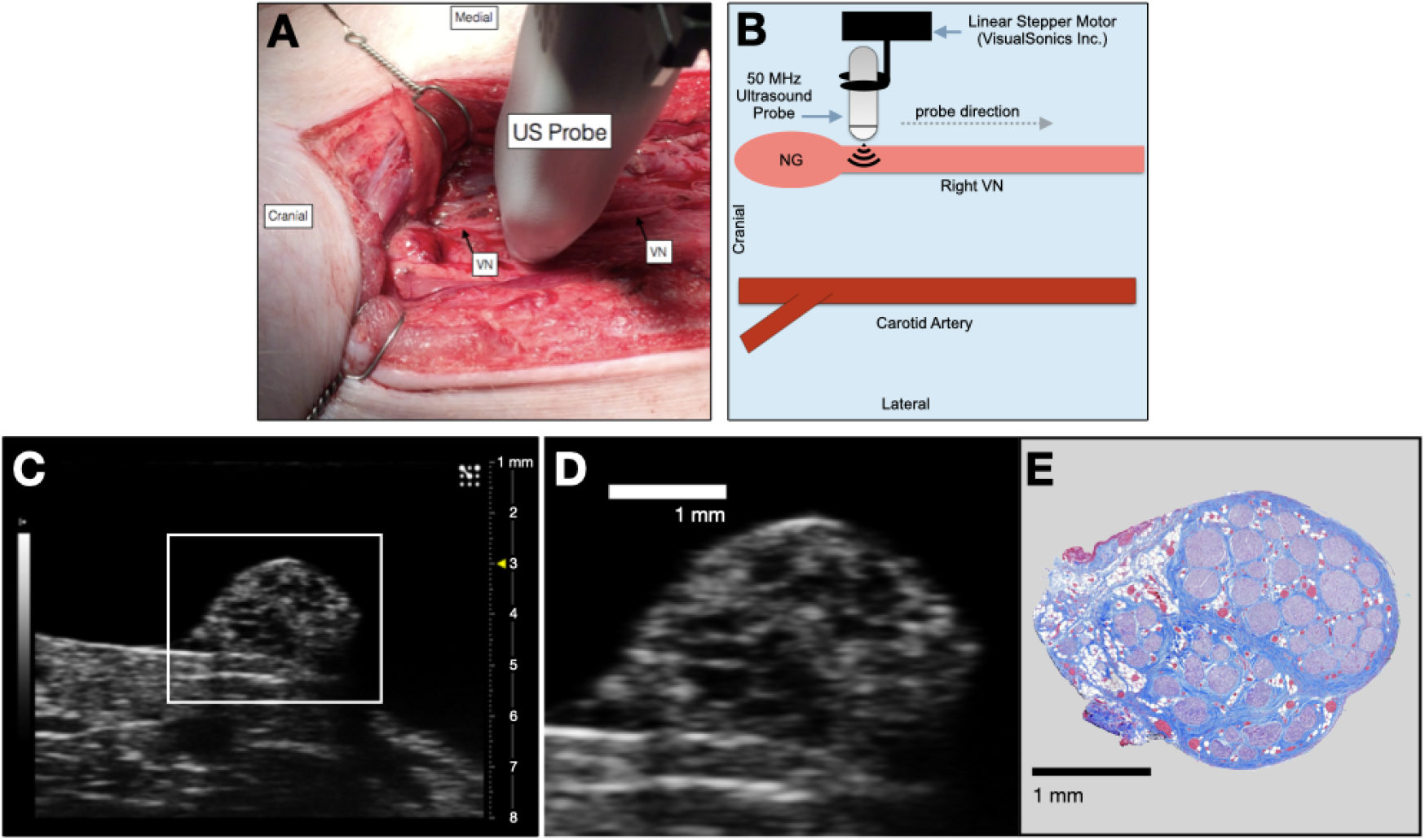
Schematic of ultrasound surgical setup in a representative subject. **(A)** The swine surgical window includes the right vagus nerve (VN) and the carotid artery, as well as the 50 MHz ultrasound probe moving in the cranial to caudal direction. Skin, muscle, and fat were retracted in this acute preparation. **(B)** Schematic of the transducer path (gray dashed arrow) as it scanned from the nodose ganglion (NG) moving caudally, approximately 10 cm. **(C)** A representative, unpaired histology section (5 μm, Gomori’s trichrome) showing the bimodal organization of the vagus nerve within the surgical window. **(D)** Ultrasound still demonstrating bimodal organization within the surgical window.

### Histology and Microdissection

The VN was exposed further to identify clearly branches extending from the main trunk, including the cardiac branches, the ST which courses parallel to the VN, and the RLN bifurcation at the level of the subclavian artery. Connective tissue was removed, and histological dye was placed along the lateral and ventral edges of the vagus nerve to maintain orientation information (Bradley Products, Inc. Davidson Marking System, Bloomington, MN).

The VN was then excised from just cranial to the nodose ganglion to the RLN bifurcation. The vagus nerves were placed in 10% neutral buffered formalin for approximately 24 hours at 4°C. Samples were then placed in a Research and Manufacturing Paraffin Tissue Processor (RMC Ventana Renaissance PTP 1530, Ventana Medical Systems, Oro Valley, AZ), and they underwent a series of standard processing steps to dehydrate, clear, and infiltrate with paraffin wax (see Settell et al. 2020 for details). Embedded samples were sectioned at 5 μm, mounted on charged slides, and stained using Gomori’s trichrome. Slides were imaged at 20x using a Motic Slide Scanner (Motic North America, Richmond, British Columbia) (Figure 1E).

### Ultrasound Video Analysis

An initial standard set of contrast optimization steps were applied to each ultrasound video, and then optimized individually to visualize the fascicular organization of each nerve. Blender (*a 3D modeling and rendering package*, Stichting Blender Foundation, Amsterdam) was used for processing and analyses, and creation of visualizations. To improve visualization of the ultrasound video and aid in fascicle organization for histological comparison, adjustements were made to the video brightness and contrast; specific Blender tools and parameters are fully described in the Supplemental Methods.

To compare ultrasound and histology, at least two histological slides were manually matched to the ultrasound images using morphological identifiers and used to seed a linear regression model. To obtain these ‘initial’ histological slices we used prominent morphological features that are consistent in both modalities as the initial pass for comparison. At the level of the cervical vagus nerve, the most common features across animals, obvious in both modalities, were the nodose ganglion and the branching point of the SLN. As these prominent features can span several slides and/or frames, an operator visually matched the histology sections and ultrasound frames to identify the most exact match to use as the starting point to seed the linear regression model. Key features used to make this visual assessment were nerve shape, size, orientation, organization of fascicles, and changes in fascicular structure from slide to slide.

Once these initial slides were identified and matched to the ultrasound, their linearly distributed position along the length of the nerve, known inter section distance, and number of frames in the ultrasound video were used to seed the linear regression model. The linear regression model was updated in an iterative fashion until all slice locations had been manually confirmed. The orientation of the histological slices within the video was determined based on the histological dye markings previously placed on the nerve (see Histology and Microdissection). The orientation of the ultrasound was identified using anatomical landmarks, such as the esophagus, direction of the projecting SLN, and the surrounding surgical pocket.

## Results

### Ultrasound of the vagus nerve to identify key anatomical features

The 50 MHz ultrasound transducer was placed within the surgical pocket, with approximately 1-2 mm between the transducer and the VN, and imaging was performed to acquire axial cross sections along the VN (Figure 1). Our Vevo^®^ linear rail system was used to acquire images starting at the nodose ganglion, and throughout the surgical pocket (approximately 10-12 cm in length), including at the typical VNS cuff locations, with fascicular organization identifiable throughout. The pseudo-unipolar cells of the nodose were identified via ultrasound as a single large fascicle, or large circular hypoechoic region within the nodose ganglion (Figure 2).

**Figure 2:**
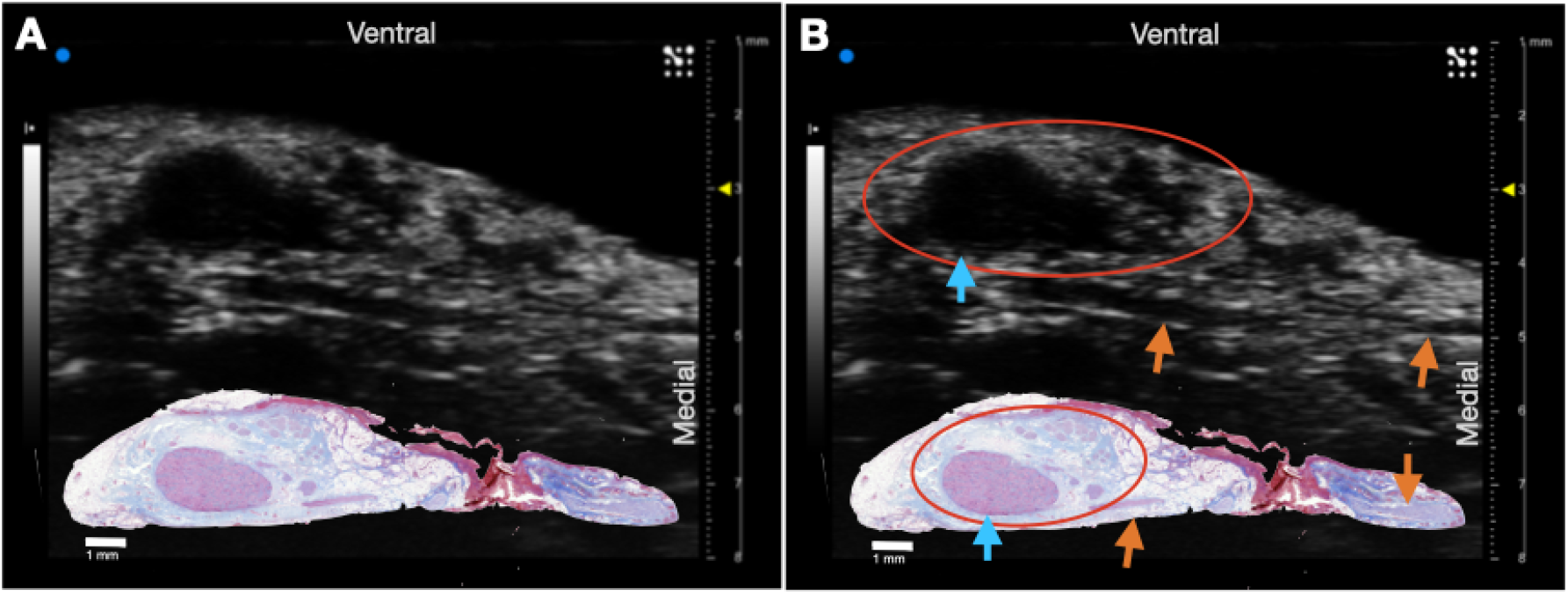
Ultrasound of the nodose ganglion in a representative subject. **(A)** Raw ultrasound and unpaired histology slice demonstrating the pseudo-unipolar cell region. **(B)** Red circles indicate the approximate boundaries of the vagus nerve, and blue arrows indicate the hypoechoic pseudo-unipolar cell region of the nodose ganglion. The superior laryngeal extends from the nodose ganglion to the esophageal cartilage with fascicles (orange arrows) running in the longitudinal direction.

In each animal, the corresponding histology was approximately matched using a combination of the regression method, fascicular organization, and identifying markers (Figure 3). The bimodal organization was visualized at various points along the length of the cervical VN (see supplemental material for the full ultrasound and histology videos, n=3). Despite visualization of fascicular structure, in some subjects the lower portion of the nerve often had a ‘shadowing’ effect as a result of the transducer being placed directly above the nerve (See Limitations).

**Figure 3:**
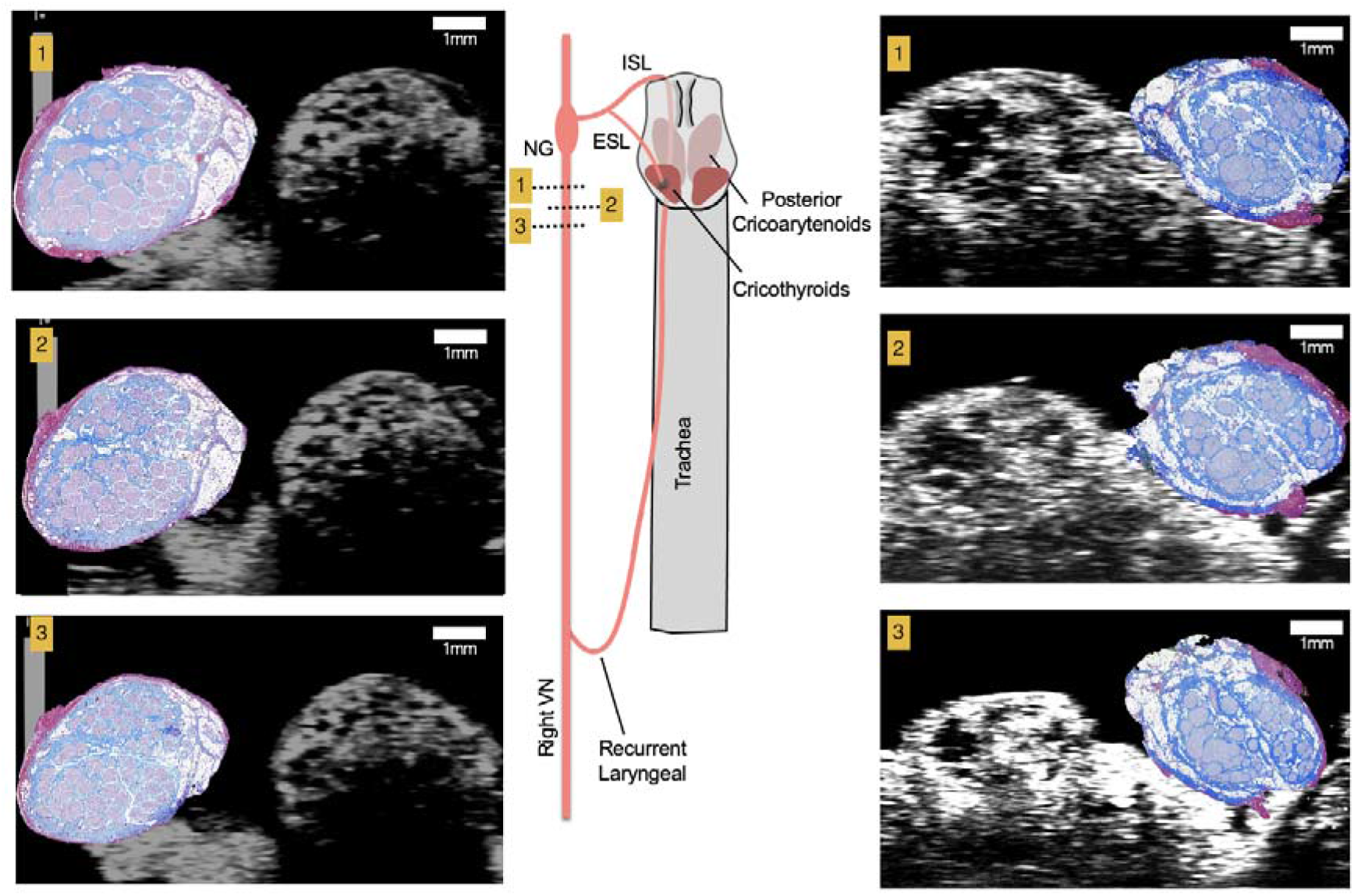
Schematic of the pig vagus nerve with corresponding locations of ultrasound and histological sections (5 μm) from two subjects (left and right). During the surgical approach the ultrasound transducer captured cross-sectional images (1 mm scale bar) along the length of the vagus nerve in the cranial to caudal direction and captured fascicular structure as shown in post-mortem histology, the histology is enlarged to show fascicular organization and is not to scale (histology insets); vagus nerve (VN), nodose ganglion (NG), internal superior laryngeal (ISL), external superior laryngeal (ESL).

To confirm the potential of the hypoechoic region of the nodose ganglion and vagotopy as a potential identifying marker in humans, we conducted microCT of a cervical VN explanted from a human cadaver (Supplemental Figure 1 and 2, Supplemental Methods). MicroCT not only confirmed the visualization of the nodose ganglion, but the extensive change of fascicular organization over a small region of the VN, highlighting the need for intraoperative visualization. Identification of fascicular organization was confirmed via histology and trichrome staining (Supplemental Figure 2B).

### Ultrasound of the superior and recurrent laryngeal branches

The RLN and SLN are somatic branches of the VN implicated in off-target activation of the deep neck muscles that produce therapy-limiting side effects (Nicolai et al., 2020). We assessed whether ultrasound could be used during the surgical procedure to visualize these branches—which are smaller in diameter than the compound VN—as this could be important to avoid off-target effects through personalized cuff placement and to inform anatomically accurate computational models of activation.

Within the surgical window the SLN and RLN were both identified using ultrasound, with visualization of fascicular structure. The SLN extends ventro-medially from the NG to muscles overlying the thyroid cartilage (Figure 4A) (Hayes et al., 2013; Settell et al., 2020). The RLN was identified as running parallel to the vagus nerve along the esophagus and inserting into the cricoarytenoid muscle. It contained far fewer fascicles, but was clearly visible (Figure 4B).

**Figure 4:**
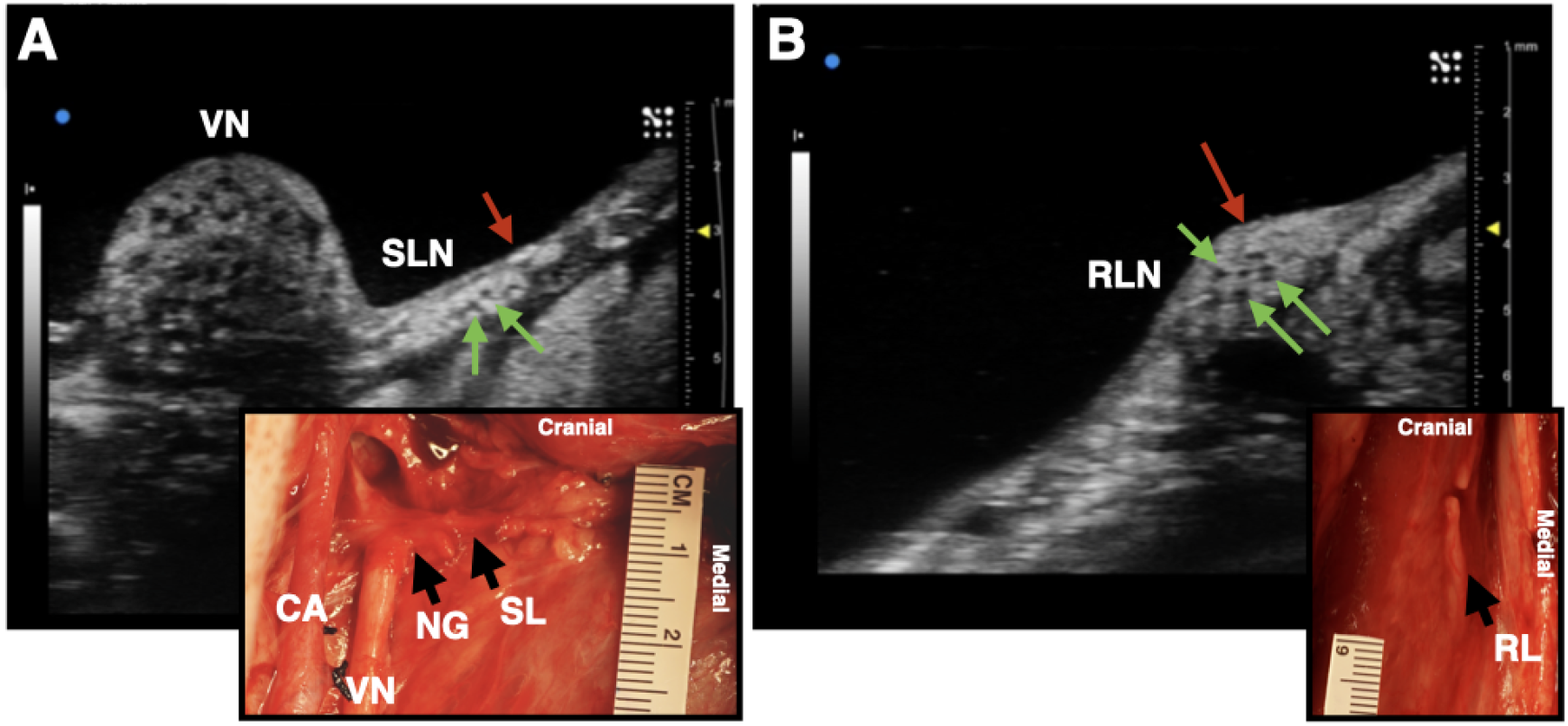
Ultrasound images of the SLN and RLN branches of the vagus nerve. **(A)** The SLN (red arrow) branching ventromedially off of the nodose ganglion (NG). Green arrows indicate fascicles within the nerve. **(B)** The RLN (red arrow), running along the esophageal groove. Green arrows indicate fascicles. Photograph insets in both **(A)** and **(B)** depict the corresponding ultrasound region: CA, carotid artery; VN, vagus nerve; SLN, superior laryngeal nerve; RLN, recurrent laryngeal nerve; NG, nodose ganglion.

## Discussion

### Improving intraoperative procedure

Surgical implantation of VNS devices has limited patient specificity (Reid, 1990; R. Terry et al., 1990; R. S. Terry et al., 1991). Briefly, the carotid sheath is located medial to the muscle and undergoes blunt dissection and is opened approximately 7 cm to expose the carotid artery, internal jugular vein, and VN. Vessel loops are used to suspend the VN while 3 cm of the nerve are dissected from any surrounding tissue to allow for proper placement of cuff electrodes. Three helical cuffs are then placed around the nerve (two stimulating electrodes and an anchor) (Giordano et al., 2017).

The simple and widely deployable introduction of ultrasound into this VNS implantation process could significantly aid in identifying 1) branches extending from the VN and implicated in producing side effects, 2) fascicular organization of the VN, 3) optimized locations for cuff placement. To validate the concept, we placed an ultrasound transducer in the surgical pocket of anesthetized pigs that were undergoing VNS experiments. The skin incision in the pig model (10-12 cm) is slightly larger than that of the human preparation (~7 cm), and the skin, fat, and muscle were retracted in the animal model to optimize transducer placement. The cavity was also filled with mineral oil to improve coupling to the nerve; however, saline was also used as a clinically translatable solution with similar results (data not shown). The VN was visible in the ultrasound with clear, identifying, features. From the ultrasound images, we visualized the fascicular organization with sufficient resolution to identify the bimodal organization (Figure 1D, Figure 3), pseudo-unipolar region (Figure 2), as well as the SLN and RLN branches (Figure 4). When these images were compared to post-mortem histology, it was determined that this approach is not only easily deployable during the procedure but captures the anatomical organization in real-time.

Using anatomical landmarks, ultrasound is effective for clinical evaluation of somatic peripheral nerves (Lawande et al., 2014) and has greater sensitivity for detection of neuropathologies than MRI (Zaidman et al., 2013). In normal, healthy peripheral nerves, the transverse section has a honeycomb-like appearance with hypoechoic areas—at the locations of fascicles—separated by hyperechoic septae. The median nerve can be consistently visualized from the mid-upper arm to the wrist using high frequency, linear-array transducers (Brown et al., 2016). Post-mortem visualization of the RLN ultrasound is used in studying neuropathologies such as vocal cord paralysis (Solbiati et al., 1985). Ultrasound has also been used clinically for detection of pathologies in peripheral nerves such as tumors and leprosy (Martinoli et al., 2000).

### Avoiding off-target effects by identifying off-target nerves

The SLN and RLN are implicated in many of the off-target effects of VNS (Nicolai et al., 2020). We aimed to evaluate the utility of ultrasound as a tool for visualizing the SLN and RLN within the surgical pocket, and identify fasicular organization. As compared to the pig model, the human SLN—which branches at the level of the nodose—may be more difficult to discern, as the nodose ganglion is typically cranial to the surgical window, and therefore simply tracing the vagus nerve back to its point of origination is not feasible. Though the SLN is smaller and contains fewer fascicles than the vagal trunk, ultrasound could potentially be used as a quick confirmation for identifying the nerve within the surgical window (Figure 4A), and for seeding computational models to inform off-target activation. As the SLN innervates several muscles of the neck that are implicated in side effects of VNS (Nicolai et al., 2020; Yoo et al., 2013), it is imperative that intraoperative placement of the VNS cuff not be in a region where current escape could activate the SLN resulting in off-target activation.

The anatomy of the SLN can vary between patients (Whitfield et al., 2010). Injuries to the external branch of the superior laryngeal (ESL) nerve, which innervates the cricothyroid muscle, result in voice changes, a common side effect of VNS (Whitfield et al., 2010). The classic anatomy of the ESL, and its relationship to traditional landmarks such as the superior thyroid artery or superior pole of the thyroid, is highly variable (Whitfield et al., 2010). Before placing the VNS cuff, the use of ultrasound to identify the external branch of the superior laryngeal, which extends into the surgical window, could aid in minimizing some of the off-target effects that occur. Visualization of the VN, SLN, and RLN can be achieved through imaging within the surgical pocket.

Non-invasive imaging of the VN has been conducted both in patients (Park et al., 2011) and cadavers (Knappertz et al., 1998), with visualization of the carotid artery, jugular vein, and VN. However, resolution was poor and the only visually obvious components were the hypoechoic jugular vein and carotid artery, with the VN difficult to identify (Knappertz et al., 1998). There has been significant work in creating a database of ultrasound images of the VN to provide neurosurgeons with a resource for predicting the location of the VN and the distribution of the depths of the nerve from the skin’s surface (Inamura et al., 2017). Though the use of ultrasound in this manner highlights the ability to view the VN non-invasively in relation to the carotid artery and jugular vein, it also demonstrates the poor resolution for viewing fascicular strucuture, and other pertinent branches (ESL, RLN). Our study demonstrates the degree to which ultrasound information within the surgical window could be personalized, not only in terms of VN location, and fascicular organization, but the location of surrounding structures. A patient-specific surgical approach, tailored by ultrasound, would allow the surgeon to consider variations in vagal branching and location.

Along with informing electrode placement, patient-specific ultrasound images could inform computational models of VNS. Computational models are critical for the development and application of neurostimulation devices, specifically in terms of optimizing the post-surgical programming process. Individualized models, seeded by patient-specific fascicular organization obtained from ultrasound could increase the speed and process of programming, and may be critical for practically programming multi-contact electrode designs in the future. Existing models for non-invasive VNS are based on high-resolution MRI and focus solely on the activation of specific targeted fiber types (Mourdoukoutas et al., 2018). However, it has been shown that ultrasound imaging provides greater resolution and sensitivity than MRI for peripheral nerves (Zaidman et al., 2013). Future computational models should consider off-target activation for better quantitative predictions of the potential side effects of VN activation. Greater consideration must be given to the SLN and RLN in future models for VNS, which can be achieved through ultrasound visualization of vagotopy and the region surrounding the implant. Current three dimensional, MRI and finite element-based models, of compound peripheral nerves incorporate realistic geometries, as well as inhomogeneous and anisotropic electrical properties of specific nerve elements such as the perineurium and endoneurium (Mourdoukoutas et al., 2018; Pelot et al., 2018). In the future, existing finite element modeling can be used to develop more realistic VN models through consideration of VN fascicular structure, gathered from ultrasound images.

### Limitations

There are several limitations to this study that should be taken into consideration. While the pig VN is similar in size to that of the human VN (Settell et al., 2020), it is at a different depth and requires a different surgical approach. The pig surgical window contains much more fat and muscle than typical human necks and therefore requires more retraction. The retracted surgical preparation allowed for the placement of the ultrasound transducer directly above the nerve (1-2 mm), something that may need to be modified in the clinical setting.

In addition to variations in anatomy, the process of preparing the histology may cause the nerve to shrink (Stickland, 1975), which may affect our matching of the ultrasound and histology. Patterns and movements of individual fascicles as well as the general shape of the nerve were considered collectively, and thus some regions of the individual subject videos, or captured stills, may not appear to be an identical match. However, the overall appearance of fascicles in the high resolution ultrasound was sufficient enough to visualize vagotopy in terms of bimodal organization originating from the hypoechoic region of pseudo-unipolar cell bodies in the nodose ganglion.

Additionally the nodose ganglion in humans is located near the base of the skull in the jugular foramen, more cranial from the surgical window than in a pig model. However, the hypoechoic region of pseudo-unipolar cells is quite large in pigs and could potentially be identified either non-invasively (pre- or intra-operatively) or by aiming the transducer towards the ganglion. This could allow identification of the bimodal organization and subsequent tracking to the surgical window region. The feasibility of the translation of this imaging methods from human to pigs may be evaluated in cadavers.

Future ultrasound work may optimize scans based on realtime data. Image quality could potentially be improved by optimizing 1) acquisition parameters, such as contrast display settings and 2) gain and focus during acquisition to limit shadowing on the dorsal aspect of the nerve. Additionally, surgical approach may optimize images by manually scanning with the transducer versus utilizing a step-motor as in the above preparation. In this manner the orientation of the transducer can be rotated to visualize all 360° of the nerve, and minimize potential shadowing effects.

## Conclusion

Vagus nerve stimulation is FDA-approved for several indications, including epilepsy and depression, and holds promise for many other indications. However, for improved clinical VNS efficacy, fascicular organization of the VN should be considered for each patient. Ultrasound is an established method for visualization of these characteristics in somatic nerves and could be implemented during the surgical implantation of the VNS lead to inform placement of cuff electrodes and to inform patient-specific computational models.

Our findings demonstrated the ability to identify the vagotopy of the pig VN intraoperatively with a high-resolution transducer. We identified the pseudounipolar cell aggregation of the nodose ganglion and were able to visualize bimodal organization of fascicular bundles, through the cervical trunk where a VNS electrode would be placed. Our ultrasound data were paired with post-mortem histology to confirm the fascicular organization. This work highlights the potential for an intraoperative technique that could improve VNS cuff placement, aid in limiting unwanted side effects, and therefore hold promise for enabling patient-specific computational models to inform stimulation paradigms.

## Supplemental Materials

### Supplemental Methods

Briefly, brightness and contrast filters were applied to sections of video first, with brightness values ranging between 15 and 45, and contrast values between 40 and 80. If that was not sufficient for identification, a second brightness and contrast filter was applied with brightness values ranging between −15 and 25, and contrast values between 10 and 40. If the overall brightness of the video appeared to be too dark, a white balance was applied as well. The filter was placed under the brightness and contrast filter(s). For sections that required only slight adjustments, values were applied at around 0.8, for sections that were much darker a value of 0.2 was applied. In addition to contrast enhancements, a “curves adjustment” was applied to improve the image. Both x and y values ranging from 0.5 to 1 were applied, though the values presented were based on the limited sample size, and each video was assessed on a case-by-case basis.

### MicroCT of Human Vagus Nerves

In addition to the fascicular structure being identified using ultrasound, we were able to confirm this organization in humans using MicroCT (Supplemental Figure 1). Vagus nerve was harvested from a disarticulated cadaver provided by Case Western Reserve University School of Medicine. The proximal end was made cranial to the nodose ganglion and the distal cut at the clavicle. The specimen was placed in 4% paraformaldehyde fixative and stained using a standard procedure for osmium tetroxide and dehydrated for 4 days. Imaging was conducted on a PerkinElmer MicroCT imaging system (Waltham, MA) obtaining a 10 μm resolution. Scan parameters were 36 μm FOV and 36 μm reconstruction with copper and aluminum filter.

**Supplemental Figure 1:**
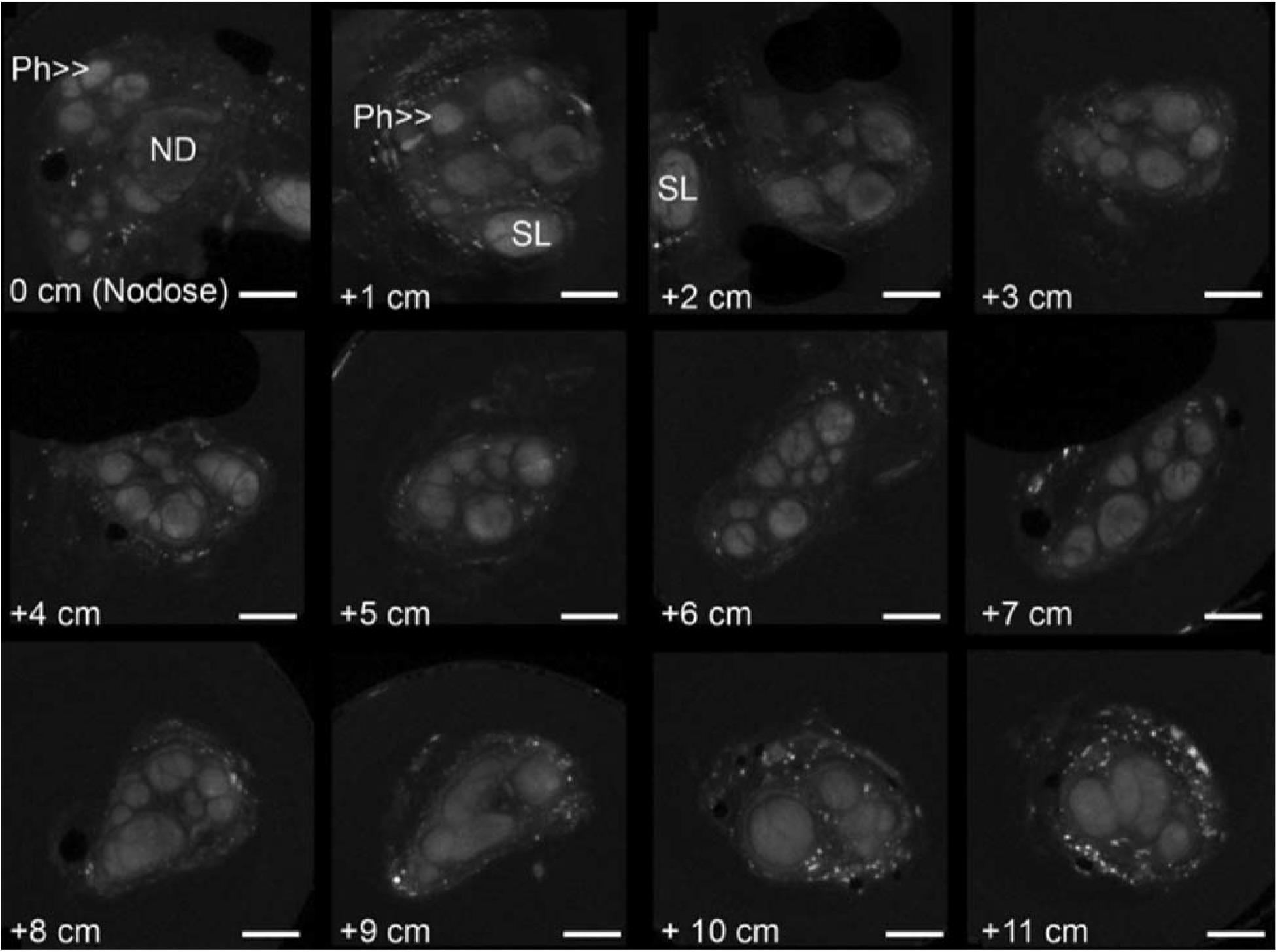
Human MicroCT of the nodose (inferior) ganglion (ND) and pharyngeal branch (Ph) (500 μm scale bar), and cervical vagus nerve. Each consecutive box (left to right) is one centimeter from the previous image indicated by the measurement in the lower left corner (distance is from 0 at the nodose ganglion).

**Supplemental Figure 2:**
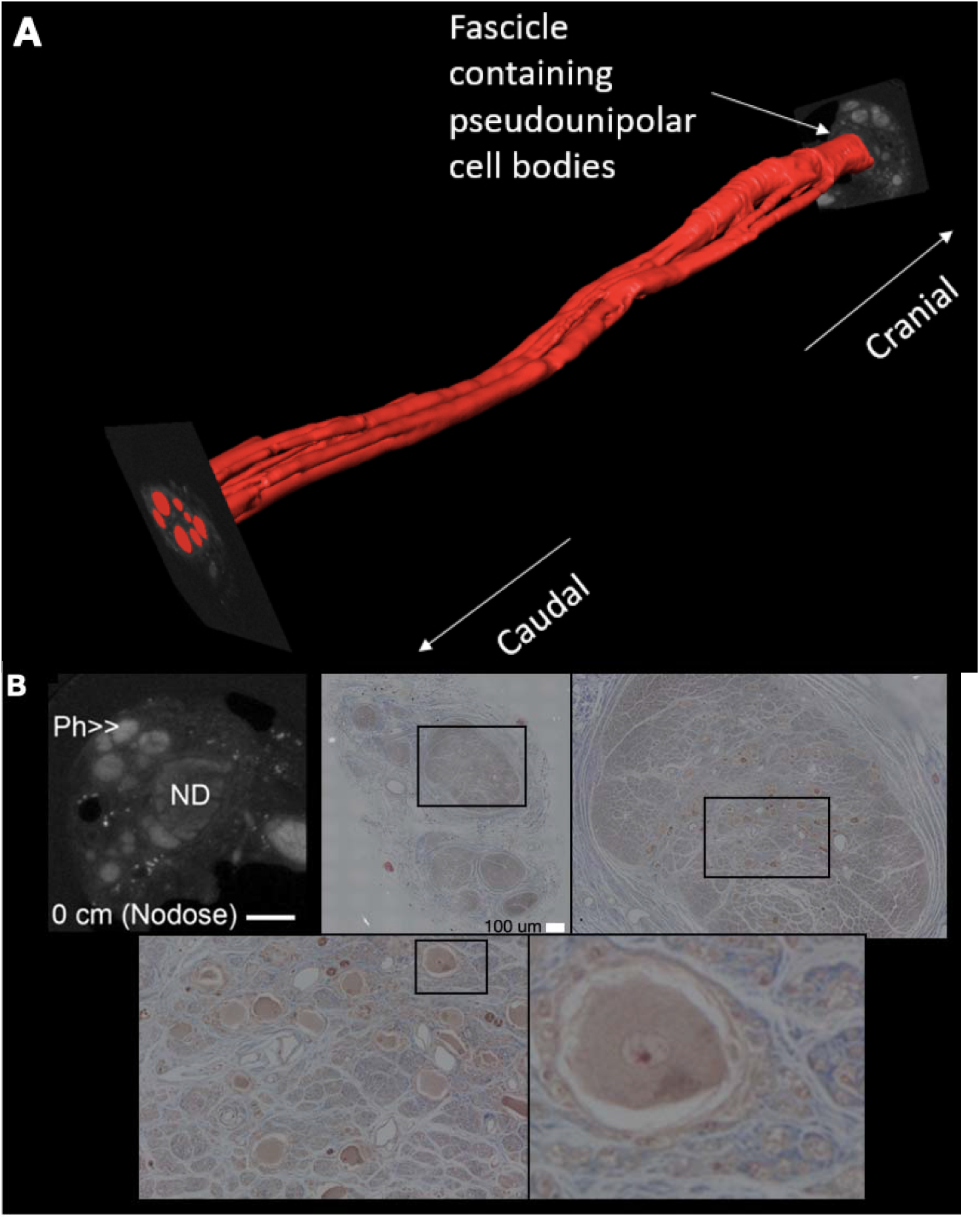
**(A)** Human vagus nerve MicroCT 3D reconstruction. The fascicles containing axons originating from the pseudo-unipolar cell bodies are highlighted in red. **(B)** Human MicroCT of the nodose (inferior) ganglion (ND) and pharyngeal branch (Ph) (500 μm scale bar) compared to corresponding histology (5 μm slice thickness, paraffin embedding, trichrome stain, 100 μm scale bar). Each histology inset depicts the next image (left to right). Definitive identification of the pseudo-unipolar cell bodies was performed by Masson’s Trichrome staining. The nodose ganglion can also be inferred from the MicroCT images based on its characteristic large relative size and appearance.

## Acknowledgements

This work was supported by the National Institutes of Health SPARC program [OT2 OD025340]. We would also like to acknowledge Dr. Jamie Van Gompel (Mayo Clinic) for his instruction on electrode placement procedures, and Andrea McConico (Mayo Clinic) for her assistance during surgical procedures.

## References

1. Boon, P., Raedt, R., Herdt, V., Wyckhuys, T., & Vonck, K. (2009). Electrical stimulation for the treatment of epilepsy. Neurotherapeutics, 6(2), 218–227. https://doi.org/10.1016/j.nurt.2008.12.003

2. Brown, J. M., Yablon, C. M., Morag, Y., Brandon, C. J., & Jacobson, J. A. (2016). US of the Peripheral Nerves of the Upper Extremity: A Landmark Approach. RadioGraphics, 36(2), 452–463. https://doi.org/10.1148/rg.2016150088

3. Cartwright, M. S., Baute, V., Caress, J. B., & Walker, F. O. (2017). Ultrahigh-frequency ultrasound of fascicles in the median nerve at the wrist. Muscle & Nerve, 56(4), 819–822. https://doi.org/10.1002/mus.25617

4. Castoro, M. A., Yoo, P. B., Hincapie, J. G., Hamann, J. J., Ruble, S. B., Wolf, P. D., & Grill, W. M. (2011). Excitation properties of the right cervical vagus nerve in adult dogs. Experimental Neurology, 227(1), 62–68. https://doi.org/10.1016/j.expneurol.2010.09.011

5. Giordano, F., Zicca, A., Barba, C., Guerrini, R., & Genitori, L. (2017). Vagus nerve stimulation: Surgical technique of implantation and revision and related morbidity. Epilepsia, 58(S1), 85–90. https://doi.org/10.1111/epi.13678

6. Giovagnorio, F., & Martinoli, C. (2001). Sonography of the Cervical Vagus Nerve: Normal Appearance and Abnormal Findings. American Journal of Roentgenology, 176(3), 745–749. https://doi.org/10.2214/ajr.176.3.1760745

7. Hammer, N., Löffler, S., Cakmak, Y. O., Ondruschka, B., Planitzer, U., Schultz, M., Winkler, D., & Weise, D. (2018). Cervical vagus nerve morphometry and vascularity in the context of nerve stimulation—A cadaveric study. Scientific Reports, 8. https://doi.org/10.1038/s41598-018-26135-8

8. Hayes, D., Nicol, K. K., Tobias, J. D., Chicoine, L. G., Duffy, V. L., Mansour, H. M., & Preston, T. J. (2013). Identification of the Nodose Ganglia and TRPV1 in Swine. Lung, 191(5), 445–447. https://doi.org/10.1007/s00408-013-9496-y

9. Howland, R. H. (2014). Vagus Nerve Stimulation. Current Behavioral Neuroscience Reports, 1(2), 64–73. https://doi.org/10.1007/s40473-014-0010-5

10. Huang, C., Lowerison, M. R., Lucien, F., Gong, P., Wang, D., Song, P., & Chen, S. (2019). Noninvasive Contrast-Free 3D Evaluation of Tumor Angiogenesis with Ultrasensitive Ultrasound Microvessel Imaging. Scientific Reports, 9(1), 4907. https://doi.org/10.1038/s41598-019-41373-0

11. Inamura, A., Nomura, S., Sadahiro, H., Imoto, H., Ishihara, H., & Suzuki, M. (2017). Topographical features of the vagal nerve at the cervical level in an aging population evaluated by ultrasound. Interdisciplinary Neurosurgery, 9, 64–67. https://doi.org/10.1016/j.inat.2017.03.006

12. Karmakar, M. K., Shariat, A. N., Pangthipampai, P., & Chen, J. (2013). High-Definition Ultrasound Imaging Defines the Paraneural Sheath and the Fascial Compartments Surrounding the Sciatic Nerve at the Popliteal Fossa: Regional Anesthesia and Pain Medicine, 38(5), 447–451. https://doi.org/10.1097/AAP.0b013e31829ffcb4

13. Knappertz, V. A., Tegeler, C. H., Hardin, S. J., & McKinney, W. M. (1998). Vagus Nerve Imaging with Ultrasound: Anatomic and in Vivo Validation. Otolaryngology–Head and Neck Surgery, 118(1), 82–85. https://doi.org/10.1016/S0194-5998(98)70379-1

14. Lawande, A. D., Warrier, S. S., & Joshi, M. S. (2014). RECENT ADVANCES IN MSK. Role of ultrasound in evaluation of peripheral nerves. Indian Journal of Radiology & Imaging, 24(3), 254–258. https://doi.org/10.4103/0971-3026.137037

15. Marciniak, C., Caldera, F., Welty, L., Lai, J., Lento, P., Feldman, E., Sered, H., Sayeed, Y., & Plastaras, C. (2013). High-Resolution Median Nerve Sonographic Measurements. Journal of Ultrasound in Medicine, 32(12), 2091–2098. https://doi.org/10.7863/ultra.32.12.2091

16. Martinoli, C., Bianchi, S., & Derchi, L. E. (2000). Ultrasonography of Peripheral Nerves. 9.

17. Morris, G. L., & Mueller, W. M. (1999). Long-term treatment with vagus nerve stimulation in patients with refractory epilepsy. Neurology, 53(8), 1731–1731. https://doi.org/10.1212/WNL.53.8.1731

18. Mourdoukoutas, A. P., Truong, D. Q., Adair, D. K., Simon, B. J., & Bikson, M. (2018). High-Resolution Multi-Scale Computational Model for Non-Invasive Cervical Vagus Nerve Stimulation. Neuromodulation: Technology at the Neural Interface, 21(3), 261–268. https://doi.org/10.1111/ner.12706

19. Nicolai, E. N., Settell, M., Knudsen, B. E., McConico, A. L., Gosink, B. A., Trevathan, J. K., Baumgart, I. W., Ross, E. K., Pelot, N. A., Grill, W. M., Gustafson, K. J., Shoffstall, A. J., Williams, J. C., & Ludwig, K. (2020). Sources of off-target effects of vagus nerve stimulation using the helical clinical lead in domestic pigs. Journal of Neural Engineering. https://doi.org/10.1088/1741-2552/ab9db8

20. Park, J. K., Jeong, S. Y., Lee, J.-H., Lim, G. C., & Chang, J. W. (2011). Variations in the Course of the Cervical Vagus Nerve on Thyroid Ultrasonography. American Journal of Neuroradiology, 32(7), 1178–1181. https://doi.org/10.3174/ajnr.A2476

21. Pelot, N. A., Behrend, C. E., & Grill, W. M. (2018). On the parameters used in finite element modeling of compound peripheral nerves. Journal of Neural Engineering, 16(1), 016007. https://doi.org/10.1088/1741-2552/aaeb0c

22. Reid, S. A. (1990). Surgical Technique for Implantation of the Neurocybernetic Prosthesis. Epilepsia, 31(s2), S38–S39. https://doi.org/10.1111/j.1528-1157.1990.tb05847.x

23. Settell, M. L., Pelot, N. A., Knudsen, B. E., Dingle, A. M., McConico, A. L., Nicolai, E. N., Trevathan, J. K., Ezzell, J. A., Ross, E. K., Gustafson, K. J., Shoffstall, A. J., Williams, J. C., Zeng, W., Poore, S. O., Populin, L. C., Suminski, A. J., Grill, W. M., & Ludwig, K. A. (2020). Functional vagotopy in the cervical vagus nerve of the domestic pig: Implications for the study of vagus nerve stimulation. Journal of Neural Engineering, 17(2), 026022. https://doi.org/10.1088/1741-2552/ab7ad4

24. Solbiati, L., De Pra, L., Ierace, T., Bellotti, E., & Derchi, L. (1985). High-resolution sonography of the recurrent laryngeal nerve: Anatomic and pathologic considerations. American Journal of Roentgenology, 145(5), 989–993. https://doi.org/10.2214/ajr.145.5.989

25. Stickland, N. C. (1975). A Detailed Analysis of the Effects of Various Fixatives on Animal Tissue with Particular Reference to Muscle Tissue. Stain Technology, 50(4), 255–264. https://doi.org/10.3109/10520297509117068

26. Taylor, T., Meer, J., & Beck, S. (2016). Ultrasound-Guided Ulnar, Median, and Radial Nerve Blocks. Emergency Medicine, 48(7), 321–324. https://doi.org/10.12788/emed.2016.0043

27. Terry, R. S., Tarver, W. B., & Zabara, J. (1991). The Implantable Neurocybernetic Prosthesis System. Pacing and Clinical Electrophysiology, 14(1), 86–93. https://doi.org/10.1111/j.1540-8159.1991.tb04052.x

28. Terry, R., Tarver, W. B., & Zabara, J. (1990). An Implantable Neurocybernetic Prosthesis System. Epilepsia, 31(s2), S33–S37. https://doi.org/10.1111/j.1528-1157.1990.tb05846.x

29. Tosato, M., Yoshida, K., Toft, E., & Struijk, J. J. (2007). Quasi-trapezoidal pulses to selectively block the activation of intrinsic laryngeal muscles during vagal nerve stimulation. Journal of Neural Engineering, 4(3), 205–212. https://doi.org/10.1088/1741-2560/4/3/005

30. Whitfield, P., Morton, R. P., & Al‐Ali, S. (2010). Surgical anatomy of the external branch of the superior laryngeal nerve. ANZ Journal of Surgery, 80(11), 813–816. https://doi.org/10.1111/j.1445-2197.2010.05440.x

31. Yoo, P. B., Lubock, N. B., Hincapie, J. G., Ruble, S. B., Hamann, J. J., & Grill, W. M. (2013). High-resolution measurement of electrically-evoked vagus nerve activity in the anesthetized dog. Journal of Neural Engineering, 10(2), 026003. https://doi.org/10.1088/1741-2560/10/2/026003

32. Zaidman, C. M., Seelig, M. J., Baker, J. C., Mackinnon, S. E., & Pestronk, A. (2013). Detection of peripheral nerve pathology: Comparison of ultrasound and MRI. Neurology, 80(18), 1634–1640. https://doi.org/10.1212/WNL.0b013e3182904f3f

